# Simulated climate change affects how biocrusts modulate infiltration and desiccation dynamics

**DOI:** 10.1101/145318

**Authors:** Angela Lafuente, Miguel Berdugo, Mónica Ladrón de Guevara, Beatriz Gozalo, Fernando T. Maestre

## Abstract

Soil surface communities dominated by mosses, lichens and cyanobacteria (biocrusts) cover most of the soil surface between vegetation patches in drylands worldwide, and are known to affect soil wetting and drying after rainfall events. While ongoing climate change is already warming and changing rainfall patterns of drylands in many regions, little is known on how these changes may affect the hydrological behaviour of biocrust-covered soils. We used eight years of continuous soil moisture and rainfall data from a climate change experiment in central Spain to explore how biocrusts modify soil water gains and losses after rainfall events under simulated changes in temperature (2.5ºC warming) and rainfall (33% reduction). Both rainfall amount and biocrust cover increased soil water gains after rainfall events, whereas experimental warming, rainfall intensity and initial soil moisture decreased them. Initial moisture, maximum temperature and biocrust cover, by means of enhancing potential evapotranspiration or soil darkening, increased the drying rates and enhanced the exponential behaviour of the drying events. Meanwhile, the warming treatment reduced the exponential behaviour of these events. The effects of climate change treatments on soil water gains and losses changed through time, with important differences between the first two years of the experiment and after five years since its setup. These effects were mainly driven by the important reductions in biocrust cover and diversity observed under warming. Our results highlight the importance of long term studies to understand soil moisture responses to ongoing climate change in drylands.

## Introduction

Arid, semi-arid and dry-subhumid ecosystems (drylands) have rainfall regimes characterized by long periods without rainfall, which also takes place as discrete pulses that largely control the activity of organisms and the rate of ecosystem processes that depend on them, including nutrient cycling, soil respiration or plant productivity, to name a few (Austin et al., 2004; Huxman et al., 2004; Reynolds, Kemp, Ogle, & Fernández, 2004). Ongoing climate change is increasing the variability of rainfall in many drylands worldwide (D’Odorico & Bhattachan, 2012; Singh & Kumar, 2015), a trend that will likely continue and be enhanced in the future (Easterling, 2000; Hughes & Diaz, 2008; Weltzin et al., 2003). Current forecasts indicate that the rainfall regime of many drylands worldwide will be characterized by a lower number of rainfall events, which will be also more concentrated in time (Dore, 2005). Thus, understanding the factors affecting water gains and losses after rainfall events is crucial not only to understand how dryland ecosystems function, but also how they are responding to ongoing climate change (Collins et al., 2014).

Biocrusts are communities formed by mosses, lichens and microorganisms (cyanobacteria, fungi, other bacteria and archaea) that live on the soil surface in drylands worldwide (Weber, Büdel, & Belnap, 2016). These communities are a prevalent biotic component in these areas, where they control the exchange of elements and energy between the atmosphere and the soil (Pointing & Belnap, 2012). Biocrusts largely affect the hydrological cycle of drylands by controlling processes such as infiltration (Belnap, 2006; Berdugo, Soliveres, & Maestre, 2014), soil water retention (Cantón, Solé-Benet, & Domingo, 2004), albedo (Rutherford et al., 2017), surface roughness (Eldridge & Rosentreter, 1999), temperature (Couradeau et al., 2016) and runoff generation (Chamizo, Cantón, Rodríguez-Caballero, Domingo, & Escudero, 2012). The hydrological impacts of biocrusts are, however, largely dependent on: i) their composition and degree of development (Chamizo, Cantón, Lázaro, & Domingo, 2013; Chamizo, Cantón, Lazaro, Sole-Benet, & Domingo, 2012) and ii) the amount, duration and intensity of rainfall events (Berdugo et al., 2014; Chamizo, Cantón, Rodríguez-Caballero, & Domingo, 2016). However, most of our knowledge on the hydrological impacts of biocrusts come from studies conducted over short periods (i.e., typical length below two years; Cantón et al., 2004; Chamizo et al., 2016; Zaady, Katra, Yizhaq, Kinast, & Ashkenazy, 2014; but see Berdugo et al., 2014). Conducting studies over multiple years is of great importance to capture inter-annual rainfall variability, which is typically very high in drylands (D’Odorico & Bhattachan, 2012), and hence to better understand how biocrusts affect soil water gains and losses after rainfall events of different amount, duration and intensity (Chamizo et al., 2016).

Climate change is impacting the biota of terrestrial ecosystems in multiple ways (Peñuelas et al., 2013), and biocrust constituents are not an exception. Increases in temperature and changes in rainfall regimes such as those forecasted for the second half of this century have been found to negatively affect the performance and cover of lichen-and moss-dominated biocrusts in South Africa (Maphangwa, Musil, Raitt, & Zedda, 2012), Spain (Escolar, Martinez, Bowker, & Maestre, 2012; Maestre et al., 2013, 2015) and the USA (Ferrenberg, Reed, & Belnap, 2015; Johnson et al., 2012; Zelikova, Housman, Grote, Neher, & Belnap, 2012). These changes are leading to shifts in the composition of biocrust communities, with reported increases in cyanobacteria at the expense of mosses and lichens (Ferrenberg et al., 2015), which underlie the dramatic changes in nitrogen (Delgado-Baquerizo et al., 2014; Reed et al., 2012) and carbon (Escolar, Maestre, & Rey, 2015; Grote, Belnap, Housman, & Sparks, 2010; Ladrón de Guevara et al., 2014; Maestre et al., 2013) cycling observed under simulated climate change in biocrust-dominated ecosystems. Despite the growing interest and literature on the impacts of climate change on both biocrusts and the ecosystem processes associated to them (Reed et al., 2016), no study so far has evaluated how climate change-induced changes in biocrust communities affect their role as modulators of water gains and losses after rainfall events. Given the important roles played by biocrusts as determinants of soil moisture after rainfall events (Berdugo et al., 2014; Chamizo, Belnap, Eldridge, Cantón, & Malam Issa, 2016), such studies are critical to better understand the hydrological impacts of ongoing climate change on dryland ecosystems worldwide.

Here we evaluate how biocrusts modulate the effect of simulated climate change on soil water gains and losses after rainfall events in a semiarid ecosystem from central Spain. For doing so we make use of an ongoing manipulative experiment that provides continuous soil moisture data for eight years in areas with different biocrust cover and under different warming and rainfall exclusion treatments (Maestre et al., 2013). This unique dataset allowed us to explore how: i) forecasted changes in rainfall and temperature may affect water gains and losses after rainfall events across multiple years with contrasting climatic conditions, and ii) simulated climate change may affect the role of biocrusts as modulators of these hydrological features. We hypothesize that climate change-induced effects on biocrust communities, being most noticeable the reduction in the cover and photosynthetic activity of dominant lichen species, already observed during the first years of the experiment (Escolar et al., 2012; Maestre et al., 2013, Ladrón de Guevara et al., 2014), will influence their capacity to affect water gains and losses after rainfall events (Berdugo et al., 2014).

## Material and Methods

### Site description

This study was conducted in the Aranjuez Experimental Station, located in central Spain (40º01’55.7’’N-3º32’48.3’’W; 590 m.a.s.l). Its climate is Mediterranean semi-arid, with average annual rainfall and temperature of 358 mm and 15 ºC, respectively (data available since 1977 from Aranjuez Meteorological Station, 40º04’N-3º32’W; 540 m.a.sl, and from an *in situ* meteorological station). The area typically registers a marked dry period from June to September with very few rains of little intensity (June, July, August and September average rain 19, 9, 9 and 22 mm respectively from 1981–2010 period. See also Figure S1). Soils are classified as Gypsiric Leptosols (IUSS Working Group WRB 2006), with pH, organic carbon and total nitrogen content values ranging between 7.2–7.7, 9–32 mg/g soil and 0.8–4 mg/g soil, respectively, depending on the microsite (open areas, vegetation and biocrust) considered (see Castillo-Monroy et al. 2010 for more details). The vegetation is dominated by *Macrochloa tenacissima* (L.) Kunth (18% of total cover), *Retama sphaerocarpa* (L) Boiss and *Helianthemun squamatum* Pers. (6% of total cover). The open areas between vascular plant patches are covered with a well-developed biocrust community that covers ~34% of the soil surface and are dominated by lichens such as *Diploschistes diacapsis* (Ach.) Lumbsch, *Squamarina lentigera* (Weber) Poelt, *Fulgensia subbracteata* (Nyl.) Poelt, *Toninia sedifolia* (Scop.) Timdal, and *Psora decipiens* (Hedw.) Hoffm. (see Maestre et al. 2013 for a full species checklist).

### Experimental design and monitoring

We established a fully factorial experimental design with three factors, each with two levels: biocrust cover (poorly developed biocrust communities with cover <25% vs. well-developed biocrust communities with cover >50%), warming (WA, control vs. 2.5ºC temperature increase), and rainfall exclusion (RE, control vs. 33% reduction in total annual rainfall). Ten replicates per combination of treatments were established, resulting in a total of 80 experimental plots. To simulate warming, we used hexagonal open top chambers (OTCs) made from methacrylate and with a size of 40 cm x 50 cm x 32 cm. To intercept rainfall, we built a 1.2 m x 1.2 m and 1 m high, metallic frame supporting three V-shaped methacrylate gutters that cover ~35% of the surface. The methacrylate gutters had a 20º inclination and on the lower side was a PVC gutter connected by a hose to a tank to collect the excluded rainfall. The warming and RE treatments were setup in July and November 2008, respectively (see Maestre et al. 2013 for additional details).

Air temperature/humidity, soil moisture, rainfall and biocrust cover have been monitored from 2009 to 2016. Soil moisture (0–5 cm depth) was continuously monitored every 2.5 hours in all treatments using replicated automated sensors (EC-5 soil moisture sensors, Decagon Devices Inc., Pullman, WA, USA). Relative air humidity, air temperature and rainfall accumulated every 10 min was also monitored using an on-site meteorological station (OnsetCorp 2.8., MA, USA). Within each plot, the total cover of the visible components of the biocrust community (lichens and mosses) were estimated at the beginning of the experiment and once a year hereafter using high resolution photographs, as indicated in Maestre et al. (2013).

### Identification and characterization of wetting and drying events

We identified wetting and drying events following the approach of Berdugo et al., (2014). We define a wetting event as any rain registered with the *in-situ* pluviometer. Rainfall events separated by at least 2.5 h without rainfall in between were considered as different wetting events. Drying events are defined as periods of ten consecutive days without any registered rain after a rainfall event. Per these criteria, we described 521 wetting and 56 drying events from February 2009 to July 2016 (Figure S1).

We used in our analyses different covariates that are important to determine soil water gains and losses after rainfall events in drylands (Berdugo et al., 2014). Four covariates were used when analyzing wetting events: initial soil moisture, rainfall amount, rainfall intensity and their interaction. Rainfall intensity was estimated as the maximum rainfall achieved in intervals of 10 minutes. We selected these covariates because they may have opposite effects on soil water gain, i.e. while rainfall amount increases soil water content, a higher intensity of the event may reduce it because it is directly related to runoff in ecosystems such as studied (Gardner, Laryea, & Unger, 1999). Initial soil moisture was estimated as the soil water content before the rainfall event. We used two covariates when analyzing drying events: the maximum temperature reached during the drying event and soil moisture at the beginning of this event, estimated as the water content after the rainfall event.

We characterized water gains as the difference between initial moisture preceding the rainfall event and the maximum moisture achieved during this event. Drying events were characterized by means of the slope and the shape of a drying curve, which describes how soil moisture decreases through time. The drying curves were obtained from the 56 drying events recorded. Each drying curve was fitted to a linear equation

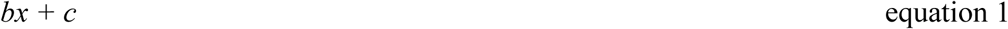

to obtain the slope (b) of the drying curve, and to a quadratic equation

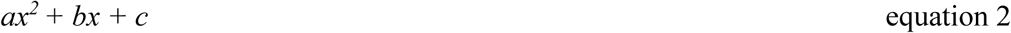

to obtain the shape (a) of the drying curve. In equation 2, *a* = 0 indicates a linear decay, *a* > 0 an exponential decay and *a* < 0 an inversed exponential decay. To improve the fitting of the drying events, we multiplied *a* by R^2^, the coefficient of determination from the quadratic regression, to obtain the shape parameter aR^2^ (Berdugo et al., 2014). The shape parameter transforms poor fitted drying curves to an aR^2^ value of 0, as these curves are considered artefacts of the statistical method and have no biological meaning.

### Statistical analyses

Generalized linear models were used to analyze the influence of the experimental treatments on the wetting and drying events. In the case of wetting events, we used the model:

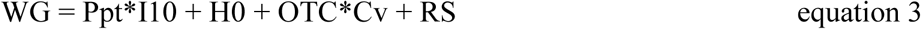

where WG is the maximum water gain at 0–5 cm depth, Ppt is the summation of rainfall amount during the event, I10 is the rainfall intensity, H0 the soil moisture preceding the rain, OTC is the effect of the open top chambers, Cv is the cover of visible biocrust components (measured from the photograph taken closest to the date of the event) and RS is the effect of the rainfall shelter. We introduced the interaction between warming and biocrust cover because we know from previous studies that warming has negatively affected the cover of biocrusts in our experiment (Escolar et al. 2012, Maestre et al. 2013). In the case of desiccation events, we built two models for the two parameters of the drying curve:

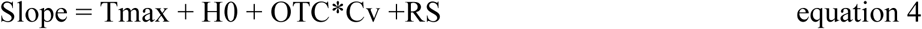

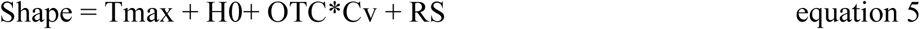

where Tmax is the maximum temperature achieved during the drying event, H0 is the initial soil moisture of this event, and the other terms are those used in equation 3.

Since biocrusts tend to colonize areas with initial low cover through time (Maestre et al., 2015), and to further investigate whether the effects of the treatments on the wetting and drying events changed through time, we repeated the analyses described above for three periods: 1–2 years, 2–5 years and 5–8 years after the setup of our experiment. We selected these periods because they showed contrasted patterns of biocrusts cover under the simulated climate change treatments of our study (Figure 1). We performed all statistical analyses using the “lm” function of the R 3.3.2 statistical software (R Development Core Team, 2011).

**Figure 1.**
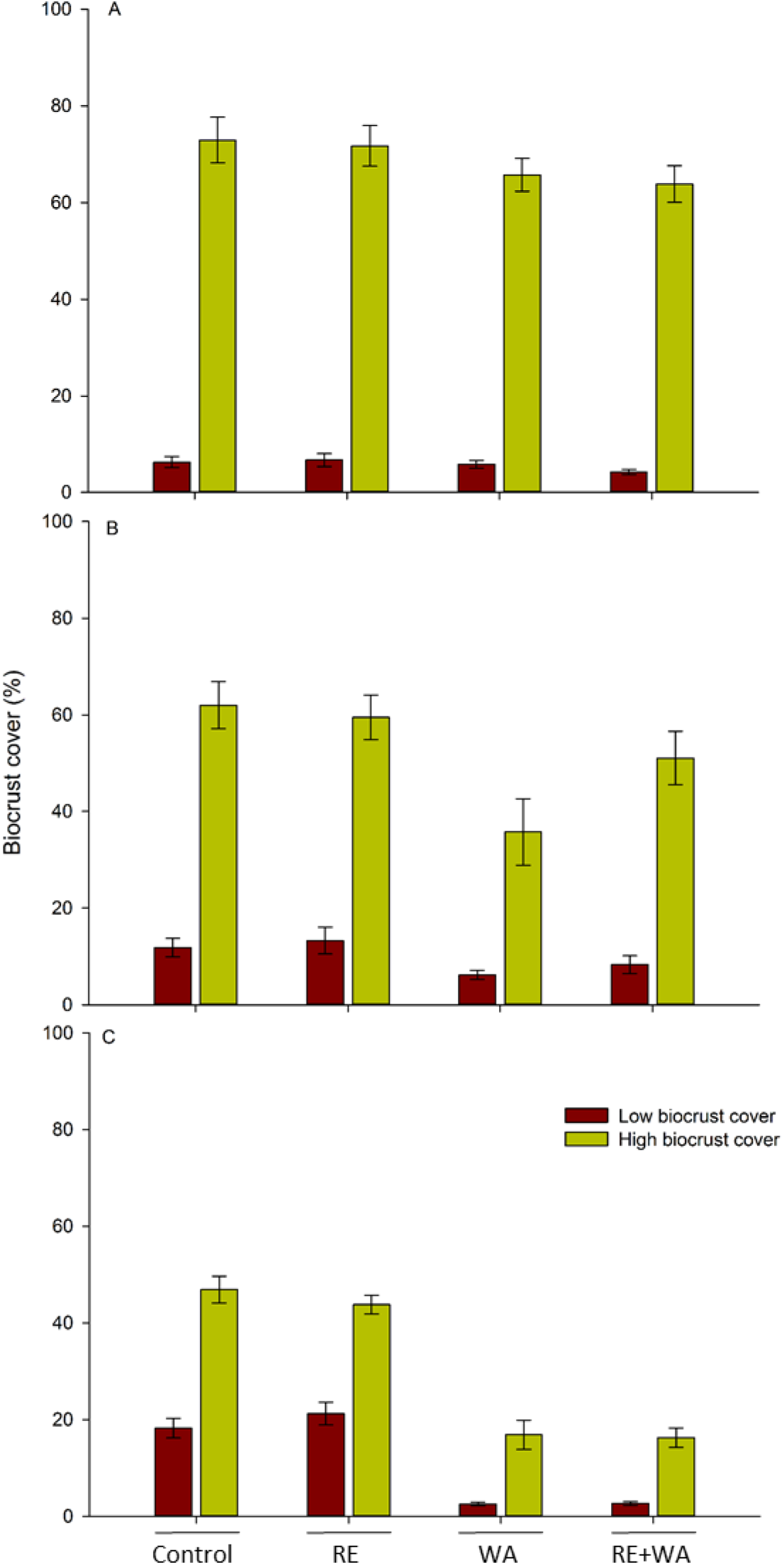
Cover of biocrusts in the different treatments throughout the duration of the experiment: 0–2 years (A), 2–5 years (B), and 5–8 years (C). WA = Warming and RE = Rainfall exclusion. Data are means ± SE (n = 10).

## Results

Soil moisture content throughout the study period accurately matched rainfall events (Figure S1). Simulated climate change treatments increased soil moisture content during the first years of the study. However, this response was inverted during the last years of the study, as both warming and rainfall reduction reduced soil moisture, particularly in areas with initially low biocrust cover.

Climatic covariates (H0, Ppt, I10 and interaction PptxI10 and Tmax) were major drivers of both the maximum water gain during wetting events and the slope and shape of the drying curve (Table 1; Figure 2). During wetting events, rainfall amount increased water gain. However, rainfall intensity decreased soil water gain during these events, even in the case of those with higher rainfall amounts (Table 1). Initial soil moisture negatively influenced water gains during rainfall events. During the drying events, initial moisture and maximum temperature increased both drying rates and the exponential behavior of the drying curve (Table 1; Figure 2 b, c).

**Table 1.** Estimates of standardized effects of covariates introduced to model water gains and losses (slope and shape of the drying curve) after rainfall events. Tmax: Maximum temperature during desiccation event; H0: initial soil moisture before the considered rainfall event; RS: rainfall shelter; OTC: warming treatment; Cv: biocrusts cover; Ppt: summation of rainfall during the wetting event; I10: rainfall intensity; F: Fisher statistic; d.f.: degrees of freedom.

| Predictor | Water Gain | Slope | Shape |
| --- | --- | --- | --- |
| H0 | -0.12 *** | -0.51 *** | 0.05 |
| Tmax | - | -0.43 *** | 0.27 *** |
| Ppt | 0.61 *** | - | - |
| I10 | -0.09 *** | - | - |
| Ppt x I10 | -0.22 *** | - | - |
| OTC | -0.10 *** | -0.06 | -0.35 *** |
| RS | 0.04 * | -0.10 | -0.02 |
| Cv | 0.08 *** | -0.19 *** | 0.07 |
| OTC x Cv | -0.06 *** | 0.09 | -0.06 |
| Adjusted R2 | 0.3945 | 0.2929 | 0.1060 |
| F | 810.7 | 55.05 | 16.47 |
| d.f. | 8/9934 | 6/777 | 6/777 |

\*\*\* = $p < 0.001$ ; \* = $p < 0.05$

**Figure 2.**
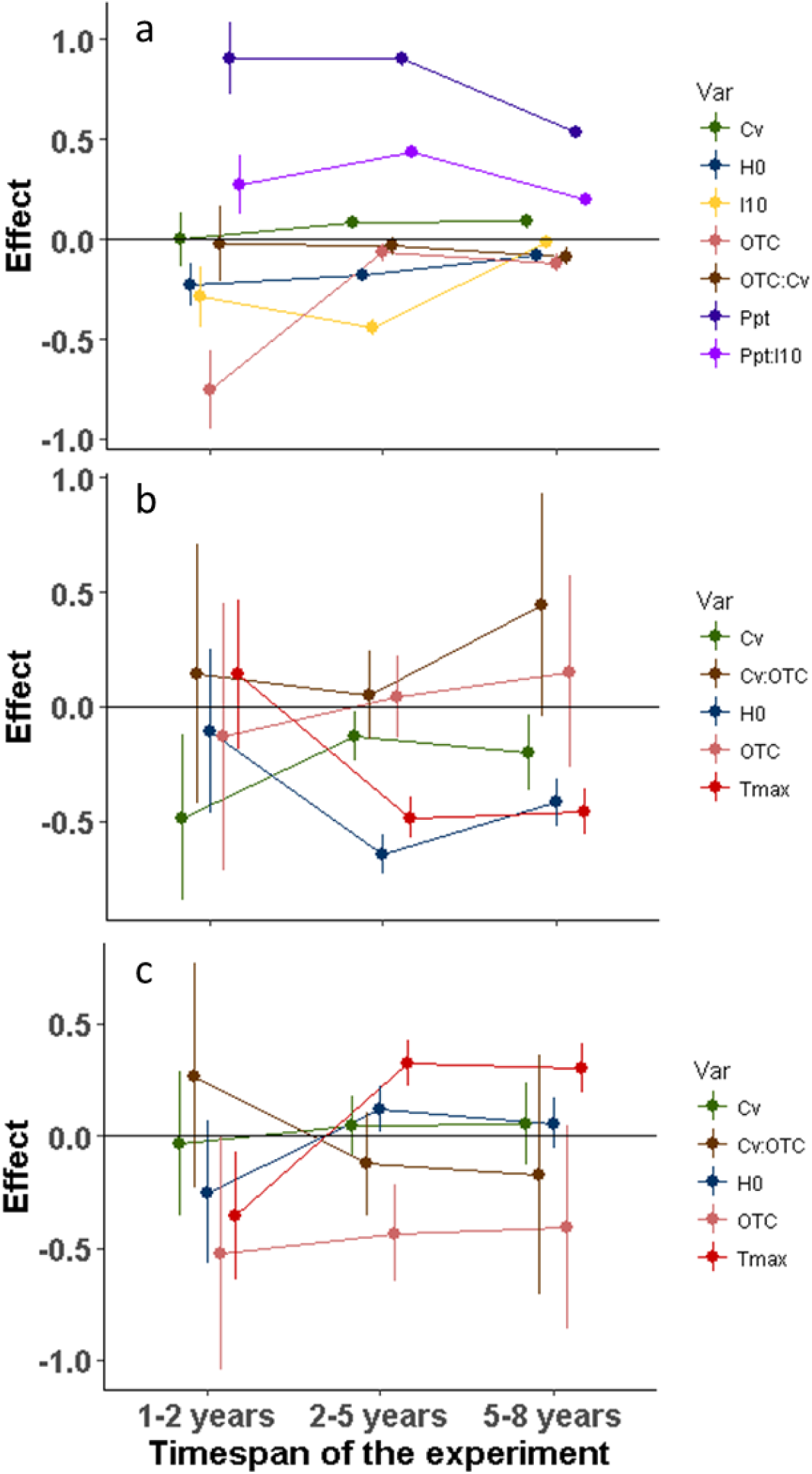
Standardized effect size of variables involved in predicting water gains (a), and the slope (b) and shape (c) of the desiccation curve after rainfall events throughout the duration of the experiment. Vertical lines represent 95% confidence intervals. Tmax: Maximum temperature during desiccation event; H0: initial soil moisture before the rainfall event; OTC: warming treatment; Cv: cover of biocrust; Ppt: summation of rainfall during the wetting event; I10: rainfall intensity.

Biocrust cover increased both water gains after rainfall events and the slope of the drying curve (Table 1). Rainfall exclusion had a positive effect only on water gains. Warming decreased water gains, but also substantially reduced the exponential behavior of the drying curve. These effects of warming were mainly evident during the first years of the experiment, and their magnitude decreased through time (Table 1; Figure 2). We also found an interaction between warming and biocrust cover when analyzing water gains, as the effect of biocrusts cover on this variable was reduced under warming (Table 1).

The effects of the different treatments on water gains and losses varied through time. As noted above, the effects of warming were particularly important and negative during the first years of the experiment (Figure 2 a). Biocrust cover negatively affected the slope of the drying curves along the experiment (Figure 2b, c). In addition, the effects of initial moisture and maximum temperature varied throughout the experiment. The effects of both initial moisture and maximum temperature on the slope of the drying events shifted to negative after the first two years of the experiment. The effect of these covariates on the shape of the drying events was the opposite. The effect of warming on the response variables evaluated was similar throughout the experiment.

## Discussion

Our analyses of an 8-yr dataset showed a clear effect of simulated climate change on soil water gains and losses after rainfall events in a biocrust-dominated semiarid ecosystem. A 2.5ºC warming reduced water gains after rainfall events, and affected the way the soil dried up after them by reducing both slope and exponential behavior of the drying curve.

An important result of our study is that the effects of treatments on soil water gains and losses changed through time, with important differences between the first two years of the experiment and after five years since its setup. These effects were mainly driven by the important reductions in biocrust cover and diversity under warming (Escolar et al., 2012), similarly to what has been reported in other climate change experiments conducted with biocrusts (Couradeau et al., 2016; Ferrenberg et al., 2015; Garcia-Pichel, Loza, Marusenko, Mateo, & Potrafka, 2013). Our findings thus emphasize the importance of conducting long-term experiments to accurately characterize potential responses to climate change in hydrological variables such as water gains and losses after rainfall events.

Soil water gains were dependent on both the amount and intensity of rainfall events. However, there was an interaction between both precipitation attributes. The greater the rainfall event, the larger the soil water gains. Nevertheless, this effect was modulated by rainfall intensity; despite its magnitude, if the intensity of a rainfall event was high, soil water gains turned to be lower than expected. Many studies have evaluated the effects of the amount and intensity of rainfall on infiltration and runoff (e.g., Cantón et al., 2004; Chamizo et al., 2012; Kidron & Yair, 1997). Our results are similar to those of Chamizo et al., (2012) and Kidron & Yair (1997), who observed that in arid and semi-arid regions under low intensity rainfall events, biocrust-dominated soils had the ability to reduce runoff when compared to bare ground soils. However, when the intensity of the rainfall event was high biocrusts lost their capacity to reduce runoff. Moreover, the higher the initial soil moisture, the lower the water gains after rainfall events. Fine textured soils show numerous cracks when are dry, which can considerably increase water infiltration. However, these cracks seal when they are wetted, and consequently runoff is favored under these conditions (Chamizo et al., 2012). In addition, wetted soils present a higher total pore volume filled with water, which reduces the capacity of the soil to further gain water (Gardner et al., 1999). Soil moisture just after the precipitation pulse, combined with the maximal temperature registered during the drying event, affected soil losses through desiccation. These results were similar to those found by Berdugo et al. (2014).

The dynamics of soil wetting and drying after rainfall events was impacted by our climate change treatments, especially by warming. Increases in the mean annual temperature imposed by this treatment (average increase in temperature ~2.5ºC throughout the experiment) led to drier soils (average reduction in soil moisture ~1% throughout the experiment), a response that also has been observed in experimental studies conducted in grasslands (Liancourt et al., 2012) and shrublands (León-Sánchez, Nicolás, Nortes, Maestre, & Querejeta, 2016) from dryland areas. Our warming treatment decreased the observed water gains and reduced the exponentiality of the desiccation curve, meaning that soil under warming gained less water and desiccated more gradually than in other treatments. These findings mimic those of Liancourt et al. (2012), who found that OTCs like those employed in our study significantly reduced soil moisture and decreased soil drying rates in a Mongolian grassland. These authors found that this reduction in the desiccation rate was a consequence of a reduction in wind speed due to the warming structures. However, we believe that the reduction in the desiccation rate observed in our case was due to a reduced water gain that leaded to a lower initial soil moisture (our OTCs were elevated 5 cm from the soil, hence allowed ventilation), and thus to a more progressive drying (reduction of the exponentiality) instead of the abrupt drying observed in a more exponential curve.

Our rainfall exclusion treatment increased soil water gains after rainfall events. This result was unexpected because this treatment effectively excluded over 33% of incoming rainfall (Maestre et al. 2013). We believe that these results may be due to the fact that we only placed one sensor under each rainfall exclusion plot, which occupies an area of 2.64 m^2^. As such, the soil moisture registered could have not provided an accurate measure of the soil surface moisture under this treatment. Therefore, the results of our study regarding the effect of rainfall shelters should be treated with caution and must be confirmed by additional studies.

The effect of biocrusts on water gains and losses after rainfall events is complex but of paramount important since they cover large areas of the soil surface in drylands worldwide. A large body of literature has discussed how the presence of biocrusts may enhance or decrease infiltration and runoff (see Chamizo et al., 2016 for a recent review). Reductions in infiltration by biocrusts have been described when these organisms smooth the soil surface (Belnap et al., 2003; Eldridge, Zaady, & Shachak, 2000) and under intense rainfalls due to enhanced hydrophobicity promoted by certain biocrust constituents and to pore clogging as consequence of the production of exopolysaccharides by biocrust-forming cyanobacteria (Warren, 2003). On the other hand, biocrusts increase soil stability (Eldridge & Leys, 2003), porosity (Felde, Peth, Uteau-Puschmann, Drahorad, & Felix-Henningsen, 2014) and surface roughness (Belnap, 2006), and other studies have highlighted how biocrust-dominated soils increase water gains and reduce runoff, maintaining a higher moisture in the top cm after a rainfall event (Cantón et al., 2004; Chamizo, Belnap et al., 2016; Chamizo et al., 2013; Chamizo et al., 2012; Chamizo et al., 2016). Our results are in accordance with these studies, as we found a significant positive effect of well-developed biocrusts on soil water gains after rainfall events (Table 1). This effect is likely to be driven by the increase in soil surface roughness by dominant lichens, which reduces runoff and increases infiltration, and by the positive effects of these organisms on pore formation in the soil, which also exerts positive effects on infiltration (Bowker, Eldridge, Val, & Soliveres, 2013; Chamizo et al., 2012).

We found that well-developed biocrusts increased both the slope and exponentiality of the drying events. Soils with a well-developed biocrust community gained more water after rainfall events, but also had higher drying rates. These results mimic those from Berdugo et al. (2014), who used another mid-term dataset (6 yrs) from the same study area. Our results suggest that evaporation is higher in biocrust-dominated than in bare ground soils. We speculate that this could be consequence of the soil surface darkening by cyanobacteria, which are abundant in our study area (Cano-Díaz, Mateo, Muñoz, & Maestre, 2017), and by some lichen species (e.g. *Toninia sedifolia*) that would thus increase surface heating and enhance evaporation, and of the increase in soil surface roughness by biocrusts, which also increases the amount of soil surface than can be heated (Kidron & Tal, 2012).

Our results indicate that ongoing climate change, and warming in particular, will affect the hydrological behavior of biocrust-dominated ecosystems both directly, by increasing evaporation, and indirectly, by reducing the cover of lichen-dominated biocrusts and by altering their ability to control water gain and losses after rainfall events. Most of the hydrological studies focusing on biocrusts have a duration lower than two years (Cantón et al., 2004; Chamizo et al., 2016; Zaady et al., 2014; but see Berdugo et al., 2014). To our knowledge, no previous study focusing on the hydrological impacts of climate change and biocrusts have used a temporal series as large as that we used. The use of such a dataset provided insights on the hydrological effects of biocrusts under simulated climate change that would not have been evident had our experiment lasted for a shorter period. These findings point to the importance of conducting mid- and long-term experiments when evaluating the hydrological responses of drylands to climate change, particularly when these are mediated by organisms that are highly sensitive to changes in climatic conditions, such as biocrusts.

## Acknowledgements

We thank C. Escolar, V. Ochoa, D. Encinar and M. Navarro for maintaining the data loggers during the study period and Agencia Estatal de Meteorología (Ministry of Agriculture, Food and the Environment, Spain) for providing climatic data. This research was supported by the European Research Council (ERC) under the European Community’s Seventh Framework Programme (FP7/2007–2013)/ERC Grant agreement n° 242658 (BIOCOM) and by the Spanish Ministry of Economy and Competitiveness (BIOMOD project, CGL2013-44661-R). A. L. is supported by a FPI fellowship from the Spanish Ministry of Economy and Competitiveness (BES-2014–067831). M. B. and M. L. G. acknowledge support from the BIOCOM project. F. T. M. & B. O. acknowledge support from the European Research Council (BIODESERT project, ERC Grant agreement n° 647038).

